# Highly multiplexed oligonucleotide probe-ligation testing enables efficient extraction-free SARS-CoV-2 detection and viral genotyping

**DOI:** 10.1101/2020.06.03.130591

**Authors:** Joel J. Credle, Matthew L Robinson, Jonathan Gunn, Daniel Monaco, Brandon Sie, Alexandra Tchir, Justin Hardick, Xuwen Zheng, Kathryn Shaw-Saliba, Richard E. Rothman, Susan H. Eshleman, Andrew Pekosz, Kasper Hansen, Heba Mostafa, Martin Steinegger, H. Benjamin Larman

## Abstract

The emergence of SARS-CoV-2 has caused the current COVID-19 pandemic with catastrophic societal impact. Because many individuals shed virus for days before symptom onset, and many show mild or no symptoms, an emergent and unprecedented need exists for development and deployment of sensitive and high throughput molecular diagnostic tests. RNA-mediated oligonucleotide Annealing Selection and Ligation with next generation DNA sequencing (RASL-seq) is a highly multiplexed technology for targeted analysis of polyadenylated mRNA, which incorporates sample barcoding for massively parallel analyses. Here we present a more generalized method, capture RASL-seq (“cRASL-seq”), which enables analysis of any targeted pathogen-(and/or host-) associated RNA molecules. cRASL-seq enables highly sensitive (down to ∼1-100 pfu/ml or cfu/ml) and highly multiplexed (up to ∼10,000 target sequences) detection of pathogens. Importantly, cRASL-seq analysis of COVID-19 patient nasopharyngeal (NP) swab specimens does not involve nucleic acid extraction or reverse transcription, steps that have caused testing bottlenecks associated with other assays. Our simplified workflow additionally enables the direct and efficient genotyping of selected, informative SARS-CoV-2 polymorphisms across the entire genome, which can be used for enhanced characterization of transmission chains at population scale and detection of viral clades with higher or lower virulence. Given its extremely low per-sample cost, simple and automatable protocol and analytics, probe panel modularity, and massive scalability, we propose that cRASL-seq testing is a powerful new surveillance technology with the potential to help mitigate the current pandemic and prevent similar public health crises.

## Introduction

Several RNA viruses have emerged in recent decades as threats to human health on a global scale (e.g. HIV, MERS, SARS, Ebola, Zika).^1, 2^ In each case, the impact of these viruses would likely have been substantially mitigated by more effective surveillance technologies and contact tracing programs. In the early stages of the COVID-19 pandemic, widely available testing and contact tracing could have blunted the explosive growth of new cases. The public health crisis has been exacerbated by lack of critical supplies in some regions, including the RNA extraction kits required for reverse transcription polymerase chain reaction (RT-PCR) based molecular testing. At the time of this writing, the United States has reported nearly 2 million confirmed cases, over 100,000 COVID-19 related deaths, and over 40 million Americans have applied for unemployment. It is widely recognized that the development of a comprehensive testing infrastructure for large-scale diagnosis and surveillance, which is rapidly reconfigurable for emerging threats, is essential to ending the current pandemic and to preventing future crises due to emerging pathogens.

Current nucleic acid tests (NATs) for SARS-CoV-2 have key limitations. Traditional RT-PCR is relatively inexpensive, but requires a separate labor and time-intensive RNA extraction step prior to nucleic acid amplification. Cartridge-based nucleic-acid tests offer rapid results and minimal sample preparation, but production of cartridges and low-throughput instruments limit scalability; further, most current testing platforms feature single pathogen targets and do not provide strain or clade-level information. Multiplexed PCR platforms and metagenomic next-generation sequencing (mNGS) technologies have demonstrated some advantages for diagnosing infections compared with more traditional approaches, but cost, low sensitivity, and complicated informatics have limited adoption for routine use.^3^ Platforms such as BioFire, Genmark ePlex and TaqMan array cards are able to identify up to 20-30 targets at a time, but their high per-sample costs (exceeding $100/test), as well as their inherently low sample throughput, severely limit their utility in the setting of the large-scale surveillance efforts.^4-7^ In the midst of the COVID-19 pandemic, innovative techniques involving targeted use of NGS have been reported, such as “SwabSeq”^8^ and “LAMP-Seq”^9^. While these methods are promising for detection of SARS-CoV-2, they may not be well suited to syndromic panel (multiplex) testing or generalized surveillance, and may provide only limited clade-level information.

Analysis of pathogen-associated RNA, versus DNA, can be valuable for several reasons. A large fraction of clinically important viruses, such as coronaviruses, have RNA genomes, and many have no DNA stage of their lifecycle. All of the viral NIAID Emerging Infectious Diseases Category A and B pathogens are RNA viruses.^10^ Further, viral mRNA can also be detected from DNA viruses that cause disease, and may provide a diagnostic advantage over DNA testing by distinguishing between active and latent infections.^11, 12^ For cellular pathogens, abundant RNA sequences, such as ribosomal RNA sequences, provide biological amplification compared to analysis of the organism’s genomic DNA, thereby enhancing detection sensitivity.^13, 14^ In addition, RNA typically degrades rapidly outside of cells, permitting differentiation between living organisms and environmental/reagent contaminants. Compared with DNA, RNA tends to be shorter and usually exists in single stranded form, making it more amenable to techniques involving probe hybridization. Finally, simultaneous analysis of viral and host mRNA expression has been shown to provide additional, clinically useful diagnostic and prognostic information about disease states.^15-17^ Importantly, the RNA analysis method presented here avoids nucleic acid extraction, which is an advantage since limited supplies of the reagents needed for this step of analysis has contributed to the disruption of large-scale SARS-CoV-2 testing efforts in the United States.^18^

Several years ago, we and others described a modified <u>R</u>NA-mediated oligonucleotide <u>A</u>nnealing, <u>S</u>election, and <u>L</u>igation with next-generation <u>seq</u>uencing (RASL-seq) assay chemistry with enhanced sensitivity. This efficient reaction utilizes the T4 RNA Ligase 2 (Rnl2), which despite being an RNA ligase, efficiently catalyzes ligation of a DNA donor probe and a chimeric acceptor probe composed of two bases of ribonucleotides at the ligation junction (**Fig. 1A**).^19-23^ In addition to the high sensitivity required for pathogen detection, RASL-seq also enables very high levels of probe set multiplexing, potentially providing the means for simultaneous analysis of pathogens, their ancestral lineages, and host immune response (**Fig. 1B**). By incorporating DNA barcodes into the primers used to amplify the ligation products, a high level of sample multiplexing is achievable, which enables very high sample throughput and extremely low per-sample cost.

**Figure 1.**
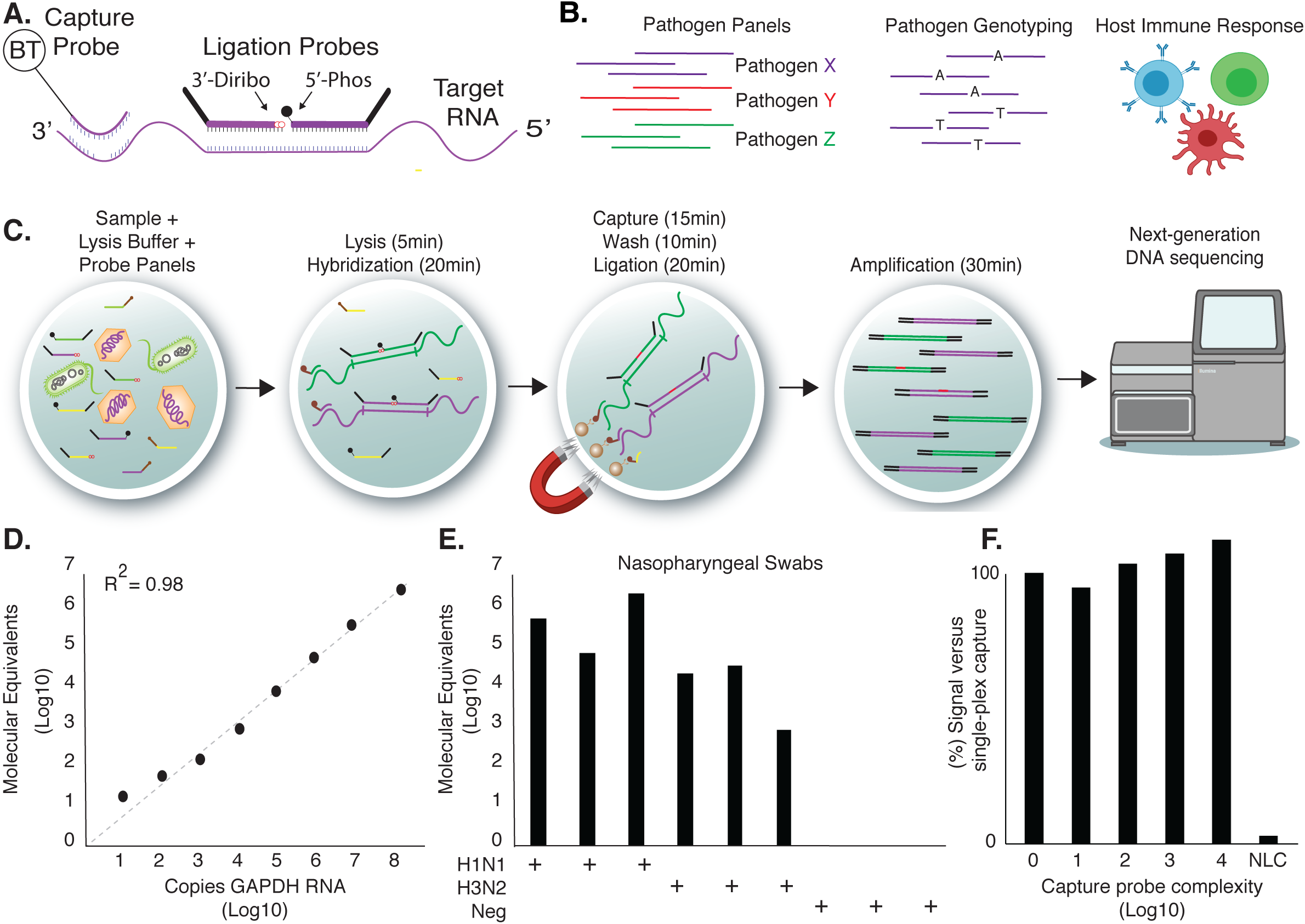
The cRASL-seq method. **A**. A ligation probe set is composed of a chimeric DNA-RNA 3’ acceptor probe and a phosphorylated 5’ donor probe. 20 nt target recognition sequences bring these probes adjacent to one another on a target RNA, enabling their enzymatic ligation. A biotinylated capture probe is included to separate the target sequence from irrelevant materials and excess ligation probes. **B**. cRASL/RASL-seq complementary assays, which can be performed in a single reaction. **C**. Sample (e.g. NP swab specimen) is added to lysis buffer containing cRASL probes. After lysis and annealing, targets are captured for subsequent ligation and sample-barcoding amplification, followed by amplicon pooling and NGS. **D**. Amount of ligation product formed on transcribed GAPDH RNA as a function of input amount; analysis by qPCR of ligation product. **E**. cRASL-seq test on a set of 9 blinded NP swabs (unextracted) from 6 patients with influenza A and 3 negative controls. **F**. Assay performed as in E, with influenza capture probe doped into a background of irrelevant capture probe at the ratio shown. For **D-F**, Molecular Equivalents are calculated by normalizing to a PCR spike-in sequence of defined copy number input.

In any probe ligation assay, it is important that excess probe be removed or destroyed in order to reduce the amount of non-specific background probe ligation.^24, 25, 26^ In contrast to previously published methods, we have incorporated the oligonucleotide-mediated capture of pathogen-associated RNA molecules, in an assay we refer to as “capture RASL-seq” or “cRASL-seq” (**Fig. 1C**). By separating targeted from untargeted RNA molecules, and thus hybridized from un-hybridized ligation probes, cRASL-seq permits extremely high assay specificity, which is especially important in the setting of diagnosing infectious diseases – particularly relevant in the early phases of an emerging pathogen threat when community prevalence is low. This method of target capture is distinct from RASL-seq analysis, which relies on immobilized oligo-dT for non-specific capture of polyadenylated mRNA.

We have previously demonstrated that libraries of ligation products are amplified with uniform efficiency,^25^ so that PCR spike-ins enable precise quantification of the copies of each ligation product formed prior to amplification. Quantification of target molecules has proven useful in clinical settings, in determining the burden of organism(s) within a clinical specimen. Furthermore, we demonstrate how cRASL-seq probes can be used for simultaneous SARS-CoV-2 detection and SNP genotyping, which has utility for tracking chains of viral transmission. Recognizing the urgent need for large-scale testing at minimal cost, we have optimized and characterized the performance of a streamlined, extraction-free protocol for direct analysis of nasopharyngeal (NP) swab specimens obtained from COVID-19 patients.

## Results

We first determined whether the standard RASL-seq assay, which does not involve RNA extraction, was compatible with analysis of COVID-19 patient mRNAs present in NP swab specimens, a matrix not previously analyzed using an oligonucleotide probe ligation technique. To this end, we utilized a large panel of RASL-seq probe sets designed to characterize human immune responses in a variety of settings (**Table S1**). A pool of 1,736 probe sets targeting 240 genes, 154 of which were assessed by analysis of exon-exon junction usage, were included in a standard RASL-seq assay with oligo-dT coated magnetic beads for capture of polyadenylated mRNA transcripts. In this experiment, an average of 727 (±46) correctly paired probe sets, corresponding to 108 (±4) genes, were sequenced at least 10 times in a given sample. We observed very high reproducibility among technical replicates (average R^2^ = 0.95 ±0.03, **Fig. S1**). High levels of housekeeping genes were detected as expected, and patterns of immunoglobulin gene expression could be reliably measured and were consistent among patients even with very different SARS-CoV-2 viral loads (determined by RT-qPCR, **Fig. S2**). These findings indicated that cRASL-seq analysis of non-polyadenylated, pathogen-associated RNA molecules might also be possible using unextracted NP swab specimens.

We determined the dynamic range of the cRASL-seq assay for detection of from 10^1^ to 10^8^ spiked-in target RNA molecules, observing exceptional linear performance over this range (**Fig. 1D**, R^2^ = 0.98). To make the cRASL-seq protocol as fast and inexpensive as possible, we performed extensive optimization to minimize the time and reagents required for each step (**Fig. 1C, Fig. S3A-C**), without compromising assay sensitivity.

We next tested the performance of the cRASL-seq assay in detecting influenza A virus in blinded, previously characterized NP swabs obtained by the Johns Hopkins Center of Excellence for Influenza Research and Surveillance (JH-CEIRS). To detect influenza A, we designed a cRASL-seq probe set targeting a conserved sequence within the M-segment, according to our previously-established design principles.^19^ A PCR spike-in standard was added at a known concentration to enable precise calculation of the ligation product copy number. Upon specimen unblinding, we observed large numbers of reads mapping onto the correctly paired M-segment probe set in all samples that contained either H1N1 or H3N2 influenza virus (**Fig. 1E**). In contrast, either zero or a small number of reads mapped to the negative control samples, providing a large signal-to-noise ratio that ranged from 10^3^ to 10^6^.

We wondered to what extent we could multiplex target capture probes, since we expected magnetic capture bead capacity to limit the level of multiplexing achievable. In order to model increasing probe pool complexity, we serially diluted the influenza M-segment biotinylated capture probe (maintained at a standard 5 pM concentration) into a background of an irrelevant biotinylated capture probe at a concentration increasing up to the binding capacity of the streptavidin coated magnetic beads used in the assay. We compared the signal of the M-segment probe in the single-plex assay (no additional capture probe) to that observed in the model multiplexed assays. We observed >90% of the single-plex signal even in a background of 10,000-fold excess irrelevant capture probe (**Fig. 1F**). While we have not explicitly tested higher levels of ligation probe multiplexing in this study, previous RASL-seq studies have employed panels of >5,000 probe sets.^27^ With appropriate design of non-interfering capture and ligation probe sets, we thus anticipate that we could achieve multiplexing of up to 10,000 probe sets, without technical artifacts.

Having established that cRASL-seq could, in principle, be leveraged into a sophisticated infectious disease diagnostics platform, we next set out to determine whether a universal protocol could be employed for diverse classes of pathogens. Important human pathogens come from all kingdoms of life. We therefore tested the streamlined, extraction-free cRASL-seq protocol for detection of the following: fungal organisms (*Candida albicans* and *Cryptococcus neoformans*) using ITS and 26S/18S rRNA (**Fig. 2A-B**); acid fast bacteria (*Mycobacterium smegmatis*) (**Fig. 2C**), gram positive bacteria (*Staphylococcus aureus*) (**Fig. 2D**), and gram negative bacteria (*Pseudomonas aeruginosa* and *Haemophilus influenzae*) using 16S rRNA (**Fig. 2E-F**); DNA virus (Human cytomegalovirus) using pp65, US34, UL5, and UL22A mRNA (**Fig. 2G**); and RNA virus (Zika virus) using genomic RNA (**Fig. 2H**). Each organism was spiked into a separate reaction in serial dilution. The combined pool of 116 probes targeting all the RNAs were tested together in each reaction (**Table S2**). Negative control reactions included full reactions without any added organism (“no template control”, NTC), as well as reactions containing the organisms but lacking the Rnl2 enzyme during the ligation step (“no ligase control”, NLC). We considered an organism detected whenever the sum of the probes’ normalized read counts was 10-fold higher than the corresponding normalized read counts from the NTC sample. In each case, we observed a strong linear correlation between normalized read counts and organism input amount across several logs of abundance, down to limits of detection that ranged from ∼1.5 to ∼150 colony or plaque forming units per milliliter **(Table 1)**. cRASL-seq can therefore be used to detect a broad range of pathogens with clinically relevant sensitivity, using a universal, nucleic acid extraction free protocol.

**Table 1.**

| Organism | Limit of detection |
| --- | --- |
| *C. albicans* | 150 cfu/mL |
| *C. neoformans* | 1.5 cfu/mL |
| *M. smegmatis* | 150 cfu/mL |
| *S. aureus* | 150 cfu/mL |
| *P. aeruginosa* | 1.5 cfu/mL |
| *H. influenzae* | 1.5 cfu/mL |
| Human cytomegalovirus | 1.5 pfu/mL |
| Zika virus | 15 pfu/mL |

**Figure 2.**
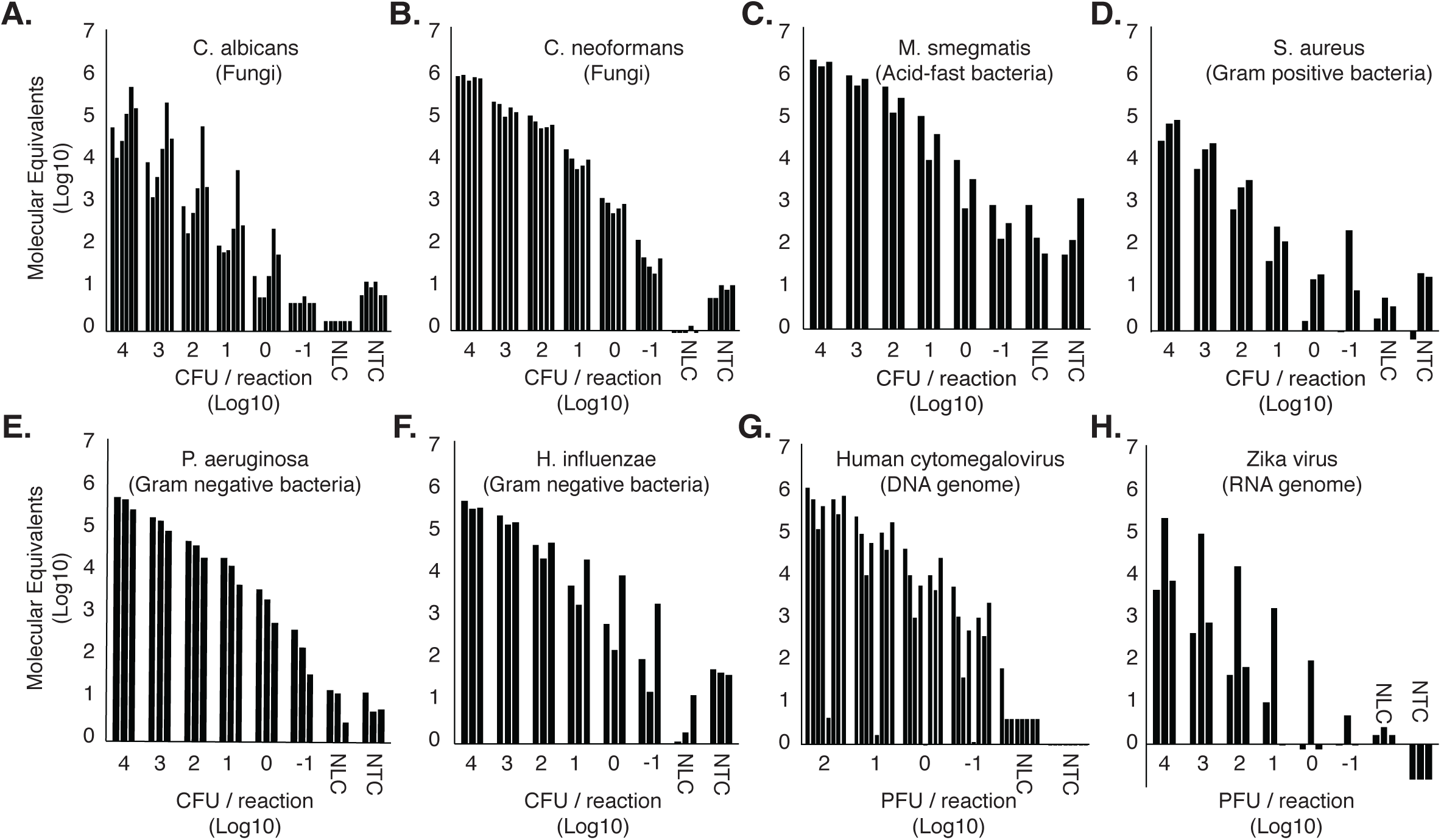
Universal cRASL-seq assay for pathogen-associated RNA analysis. Each reference organism was serially diluted into PBS and added directly to the lysis buffer and probe pool. NLC, No Ligase Control; NTC, No Template Control. The extraction free protocol of Fig. 1C was performed with all 116 probe sets in a single pool. Molecular Equivalents are calculated by normalizing read counts to a PCR spike-in sequence of known copy number. Detection at a signal >10x the NTC was used to calculate the assay’s limit of detection for each organism.

Analysis of pathogen single nucleotide polymorphisms (SNPs) has utility for distinguishing closely related organisms (e.g. human and zoonotic brucellosis), in tracing chains of viral transmission, and for detecting clade-specific differences in virulence. We therefore tested the ability of cRASL-seq probes to directly genotype SNPs from SARS-CoV-2 RNA in COVID-19 patient NP swab specimens. Genotyping probe sets were designed to share a single 3’ acceptor probe, which could pair with two alternative 5’ phospho-donor probes corresponding to the alternative genotypes (**Fig 3A, Table S3**). We placed the SNP recognition site in the center of the phospho-donor probe to maximally destabilize binding of the mismatched probe. Biotinylated capture probes were also designed to anneal within 200 nt of each genotyping probe set, to allow for some level of RNA degradation. In a proof-of-concept study, we designed genotyping probes for the 20 most entropic SARS-CoV-2 SNPs reported in the Nextstrain database^28^ (queried on 3/16/2020). These 20 SNPs span the majority of the SARS-CoV-2 genome, ranging from genomic position 241 to 29,095.

**Figure 3.**
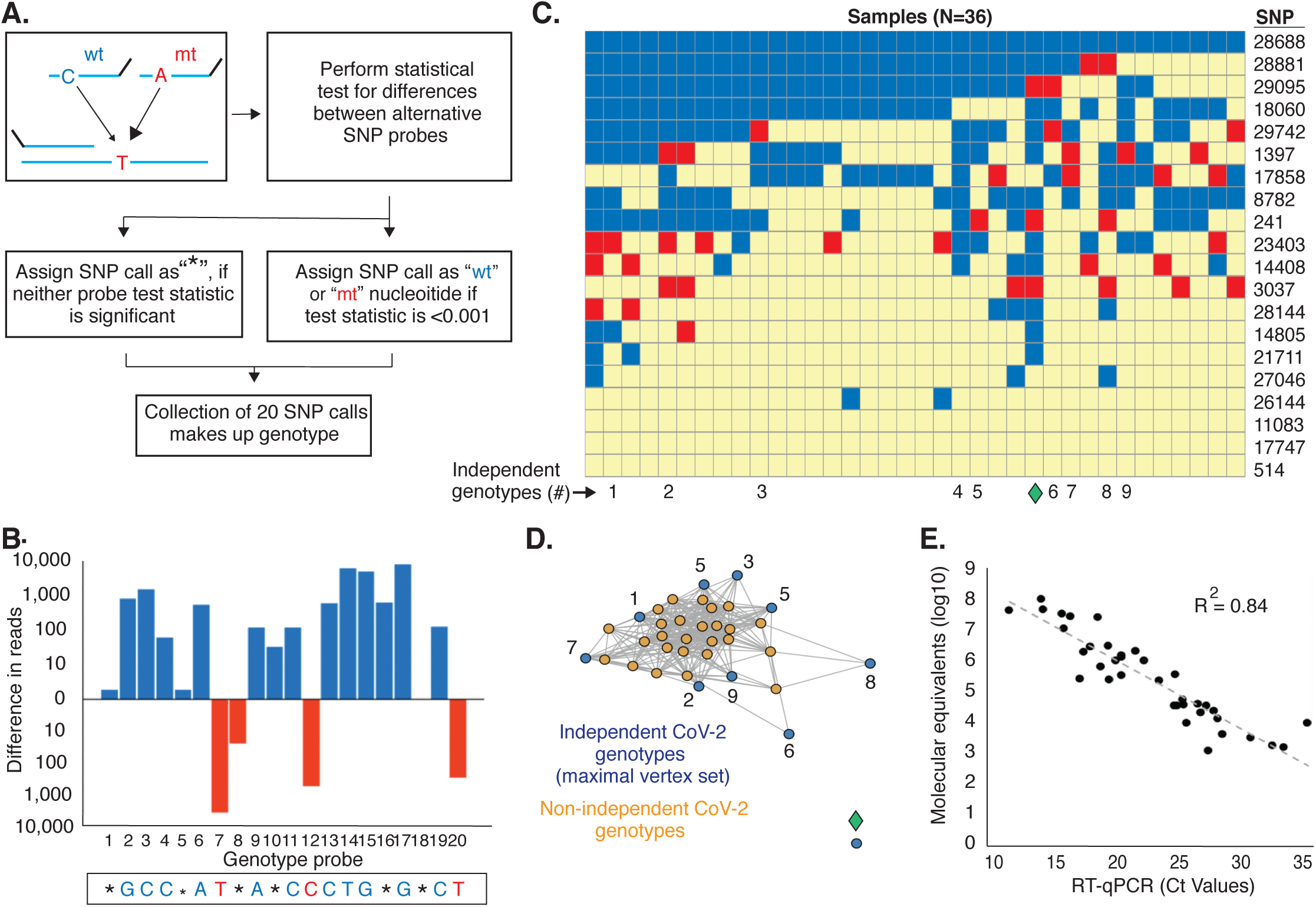
Multiplexed SNP genotyping of SARS-CoV-2 gRNA directly from unextracted NP swabs. **A**. A probe pair is designed with SNP position in the middle of the 5’ phospho donor probe. A base-calling algorithm is applied to the reads from each alternative probe. **B**. 1 of 20 positions had a base call for the reference Washington isolate, which matched perfectly against the known genotype. **C**. The 3 samples from the set of 40 PCR+ samples analyzed, which had 5 or more base calls 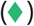. Red indicates mutant and blue indicates wildtype versus the reference Wuhan seafood market isolate. **D**. Network graph depicts each observed genotype (each individual node), two of which are linked if they do not have conflicting SNPs at any position. The blue nodes indicate a maximal vertex set of independent genotypes detected among the 3 samples that passed QC. **E**. Comparison between reads from a SNP typing cRASL-seq probe set in the N gene, versus the Ct value from the RT-qPCR (n=37, 3 samples missing Ct values).

For each SNP, the number of reads from the reference (“wildtype”) probe sequence is compared against the number of reads obtained from the non-reference (“mutant”) probe sequence. If one of the probe sets is preferentially incorporated into the ligation product (fold-difference > 1.5; p-value < 0.001, binomial test), the base is called and assigned to the position. If the probes do not have sufficient reads or they are not significantly different, no base is called and a “wildcard” is assigned to the position. The string of assigned bases and wildcards are then compared to the strings of corresponding bases from each SARS-CoV-2 genome deposited in the GISAID database (**Figs. 3A, S5**). The genotyping assay was first tested using purified reference SARS-CoV-2 gRNA. At the highest input concentration, 2×10^5^ copies per reaction, 14 of the 20 bases were called in both technical replicates, and these genotypes matched perfectly to the sequence of the isolate from which it originated (hCoV19/USA/WA1/2020|EPI_ISL_404895|20200119, **Fig. 3B**). As the input gRNA amount decreased, significantly fewer bases were called with high confidence. We assessed the reproducibility of the assay by testing 8 NP swab specimens in duplicate and comparing the results (**Fig. S4**). All technical replicates agreed with each other at a cutoff of 5 SNPs called.

We used the 20 SNP SARS-CoV-2 genotyping panel to analyze 40 NP swab specimens obtained from patients with RT-qPCR proven COVID-19. Of these, 5 or more SNPs could be called in 35 of the samples. These genotypes are displayed in **Fig. 3C**. To better understand the relationships among the observed genotypes, we used a network graph approach in which genotypes (nodes) are linked (share a connection) if they do not differ in any of the called SNPs (**Fig. 3D**). Using this network analysis, and by calculating the maximal independent vertex set, we were able to conclude that the infections among these 35 cases could be attributable to at least 9 distinct SARS-CoV-2 ancestral lineages, and thus at least that many distinct chains of local transmission. The reference Washington State isolate was unconnected to our Baltimore network. The observed genotypes could additionally be associated with geographic locations, based on their matches to sequenced isolates in the GISAID database (**Fig. S5**).

Finally, we wondered whether viral load could be simultaneously estimated using data from genotyping probe sets alone. After background subtraction using no ligase controls and seasonal coronavirus samples, a probe set targeting a SNP in the nucleocapsid gene reported positive values in all 40 PCR+ samples (100% detection sensitivity). Furthermore, **Fig. 3E** illustrates how well correlated these values are with RT-qPCR Ct values (R^2^=0.84), indicating that cRASL-seq can be used to simultaneously determine viral load and viral genotype at high sensitivity, high throughput and very low cost.

## Discussion

We have developed a generalized version of the RASL-seq technology, called “capture RASL-seq” or “cRASL-seq”, and demonstrated its utility for highly multiplexed molecular analyses of pathogens and host responses directly from NP swab specimens, using a universal and streamlined protocol that does not require up-front nucleic acid extraction or reverse transcription. The cRASL-seq protocol can easily be performed manually in a biosafety cabinet, does not require centrifugation or vortex steps that risk aerosolizing virus, does not rely on any specialized equipment, and incorporates easily scalable sample barcoding to dramatically reduce per-sample sequencing cost and increase throughput. Though not implemented here, the simple workflow can be easily automated using liquid handling instrumentation. For relatively low complexity probe panels, for example a simple SARS-CoV-2 panel, tens of thousands of sequencing reads per sample may provide sufficient sensitivity. A high output Illumina NovaSeq 6000 instrument run, which can generate up to 10^10^ single end reads, would therefore provide sufficient depth to analyze >100,000 samples at once. Another advantage of the cRASL-seq methodology is the extremely low per-probe assay concentration of 5 pM. A typical 100 nmol oligonucleotide synthesis scale will therefore yield a sufficient quantity of probe for tens of millions of 100 μl cRASL-seq tests, at a per-test probe cost below one cent. The major cost-driving components of the reaction are the ligase, polymerase and magnetic beads. Enzyme production could be scaled up to reduce costs, while less expensive streptavidin capture matrices may be adapted to replace the magnetic beads. At the scale of millions of tests, it is therefore feasible to reduce the per-test reagent and sequencing costs to below one US dollar.

Obtaining SARS-CoV-2 genotype information as part of a large-scale surveillance effort would have key benefits. Capturing viral genetics could potentially identify chains of transmission, enabling more effective contact tracing and policymaking, while also detecting and tracking emerging clades with enhanced or diminished virulence. There is also intense investigation into the role that host genetics plays in COVID-19 disease severity. As these genotypes are defined, RASL-seq probes that distinguish host alleles could be additionally incorporated into the assay. In this study, we separated the cRASL-seq analysis of SARS-CoV-2 and the RASL-seq analysis of host immune responses into two separate reactions. However, by designing non-interfering probe panels and balancing the proportion of streptavidin coated magnetic beads, versus oligo-dT coated magnetic beads, it should be straightforward to perform the two assays simultaneously in a single reaction.

The cRASL-seq methodology is not without limitations. While we have demonstrated a sensitivity comparable to single-plex RT-qPCR, the limits of detection are governed by overall sequencing depth, which can be reduced by consumption of reads from highly abundant ligation products (due to a high load of a co-infecting virus for example). However, since the ligation products are all amplified with a high degree of uniformity, simple RNA spike-in or PCR spike-in sequences can be used to determine the lower limit of detection sensitivity for each reaction. Analysis of host transcripts can also be used to assess sample acquisition sufficiency, a known source of false negative test results.^29^ Another important concern for COVID-19 molecular diagnostics is the turnaround time. When NGS is used to read out the cRASL-seq assay, the testing turnaround time is unlikely to be less than ∼24 hours with currently available instrumentation. Per-sample sequencing cost considerations will favor analysis of large sample batches, which could further increase turnaround times. For large-scale regional and national level surveillance purposes however, an occasional one to two days of self-quarantine while awaiting test results may be acceptable, given the costs and limitations associated with alternative methods. Faster, non-NGS-based readouts of cRASL probe ligation products may also be developed. For example, isothermal amplification, followed by array or test-strip hybridization may prove more applicable in the point-of-care setting. Regardless of the readout, a robust, rapidly reconfigurable, multiplexed, inexpensive and high sample throughput platform for molecular surveillance, such as the one described here, will facilitate curbing the COVID-19 pandemic and preventing future outbreaks from becoming pandemics.

## Materials and Methods

### Probe design and synthesis

For each target sequence from the reference organisms (target sequences described in Results, and Supplemental **Table S4**), we identified 40-mer sites for ligation probes using CATCH.^30^ To avoid overlapping probes, we set a stride of 40, allowing no mismatches and bypassing the cover extension in the design.py program (-pl 40 -l 40 -ps 40 -m 0 -e 0). We excluded probes that aligned against any target sequences from the other organisms, with an e-value smaller than 10^−3^, using MMseqs2.^31^ A similar design pipeline was employed for 20-mer capture probes with a final filter step to remove any overlapping ligation and capture probes. Finally, ligation and capture probes were filtered for binding properties using previously reported Primer3 conditions.^19^ The design of the immunoglobulin gene expression probe panel was reported previously.^19^ Ligation probes and capture probes (3’-diribonucleotide terminated acceptor probes, 5’ phosphorylated probes, and biotinylated capture probes) were synthesized by Integrated DNA Technologies (Coraville, IA 52241, USA). Probes were diluted in water to 100 μM, mixed in equimolar amounts to create multiplexed panels, and then aliquoted and stored at −80°C (**Table S1-S3**).

### Spike-ins and reference organisms

The synthetic PCR spike-in sequence used for determining molecular equivalents is a 74 nt oligo with a pseudo 40 nt ligated sequence flanked by the external 17 nt PCR1 primer binding sites: 5’-GGAGCTGTCGTTCACTCTGTCTCGGAGCTTACAGTrArU-TGACACTCAATCGGTCGCGTAGATCGGAAGAGCACAC-3’). The 40 nt irrelevant internal sequence is a scrambled version of a ligated GAPDH probe set. Reference organisms were purchased from American Type Culture Collection (ATCC, Manassas, VA) (**Table S4**). Organisms were reconstituted according to protocols provided by ATCC, aliquoted into single-use samples and stored at −80°C until used. The Zika virus isolate was from a patient in Cali, Colombia, and was grown in Vero-E6 cells. The infectious titer of the virus (7.1 × 10^6^ pfu/mL) was determined in the culture supernatant by a plaque assay. The full-length GAPDH ORF (RefSeq: NM_002046.6) was subcloned into a custom vector, linearized and transcribed *in vitro* using the Hi-Scribe T7 High Yield RNA Synthesis Kit (NEB, Ipswich, MA). GAPDH IVT-RNA was purified by precipitation with lithium chloride followed by column purification using the RNeasy Mini Kit (Qiagen, Hilden, DE). Purified GAPDH IVT-RNA was quantified by nanodrop, aliquoted into single-use samples and stored at −80°C until used for the dynamic range experiments.

### Nasopharyngeal swab specimens

NP swab specimens were collected from patients after informed written consent was obtained, under a protocol approved by the local governing human research protection committee. The Johns Hopkins Influenza Research and Surveillance (JH-CEIRS) program’s human subjects protocol was approved by the Johns Hopkins School of Medicine Institutional Review Board (IRB): IRB90001667 and NIH Division of Microbiology and Infectious Diseases: Protocol 15-0103. Unextracted NP swab specimens (n=9) were de-identified, blinded and provided for further analysis. Secondary use of all unextracted COVID-19 patient NP swab specimens (n = 40) was exempted by the Johns Hopkins University School of Medicine Institutional Review Board protocol (IRB00086059, and IRB00221396, **Table S4**). An RT-qPCR diagnostic test (RealStar® from Altona Diagnostics, Hamburg, Germany)^32^ was performed on all COVID-19 patient NP swab specimens used in this study. NP swab specimens were stored either at −20°C or −80°C.

### cRASL and RASL assays

Samples (39 µL) were added to 61 µL of a hybridization reaction mix containing 1X-SSC, 5 pM of each ligation and capture probe, 40 U of Protector RNase Inhibitor (Roche Diagnostics, Mannheim, Germany) and 50 µL of 2X DNA/RNA Shield (Zymo Research, Irvine CA) in a total reaction volume of 100 µL. Reactions were heated for 5 min at 95°C followed by 20 min annealing at 45°C. 5 µL of a 50/50 slurry of streptavidin coated magnetic beads (Dynabeads MyOne C1, Thermo Fisher Scientific, Waltham MA) in 1x-PBS was added to each cRASL reaction and incubated with gentle shaking for 15 min at room temperature. To each RASL reaction, 5 µL of Oligo-(dT)_25_ beads (Dynabeads, Thermo Fisher Scientific, Waltham MA) were added and incubated with gentle shaking for 15 min at 25°C. Beads were collected on a magnet for 2 minutes and washed twice with 1X-SSC buffer, followed by a final wash with 1X Rnl2 reaction buffer (50 mM Tris-HCl, 10 mM MgCl2, 5 mM DTT, 1 mM ATP, pH 7.6). 10 µL of the ligation reaction containing 30 U of Rnl2 (Enzymatics, Beverly MA) in 1X Rnl2 buffer was incubated with the beads in suspension for 20 min at 37°C. Following ligation, beads were collected for 2 min on a magnet and then resuspended in 25 µL PCR master mix containing PCR1 primers and Herculase-II (Agilent, Santa Clara CA). PCR cycling was as follows: an initial denaturing step at 95°C for 2 min, followed by 30 cycles of: 95°C for 20 s, 53.5°C for 30 s, 72°C for 30 s, with a final extension of 72°C for 3 min. Two microliters of the pre-amplification product were used as input to a 20 µL dual-indexing PCR reaction for 10 cycles with primers containing 8-mer i5 and 8-mer i7 barcodes and the P5/P7 Illumina adapters. PCR cycling was as follows: an initial denaturing step at 95°C for 2 min, followed by 10 cycles of: 95°C for 20 s, 58°C for 30 s, 72°C for 30 s, with a final extension of 72°C for 3 min.

### Library preparation and sequencing

Barcoded PCR products were analyzed individually or as a pool on a 3% agarose gel to confirm amplicon size and purity. Barcoded PCR products were pooled and purified using NucleoSpin PCR Clean-up columns (Mackery Nagel, Duren DE). Pooled libraries were sequenced on an Illumina NextSeq 500 instrument (Illumina, La Jolla CA), using a single-end 50-cycle protocol with a custom read 1 sequencing primer and custom i5/i7 sequencing primers as previously described.^17^

### Quantification of ligation products

Relative quantification of ligated products (molecular equivalents) were calculated using a synthetic PCR spike-in (described above), which was added to PCR1 reactions at 3,000 or 5,000 molecules/reaction. For samples measured by qPCR, molecular equivalents were calculated using the following equation: 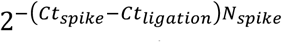 where *N*_*spike*_ is the molecules of spike-in added to PCR1 reactions. For samples analyzed by sequencing, a pseudocount was added to each probe set’s read count and the molecular equivalents were calculated by taking the ratio of ligation product read count to spike-in read count, multiplied by the molecules of spike-in added to the PCR1 reaction. For comparison of read counts with Ct values, we performed baseline subtraction on spike-in normalized probe values by subtracting the maximum normalized value of each probe from the four negative samples (two no ligase controls and two seasonal coronavirus samples). The corrected values of the best performing genotyping probe set, which targets the N gene at genome position 28,688, was plotted against clinically-determined SARS-CoV-2 RT-qPCR Ct values. We fit an exponential regression to the data, and graphed the resulting data points and regression on a semilog plot. For analysis of host gene expression, psuedocounted and GAPDH-normalized values were obtained for each probe set. For the analysis of Ig gene expression, the normalized values of all probe sets corresponding to each Ig (3 probes each for IGHA, IGHD, IGHE, and IGHM, and 8 probes for IGHG1-4) were summed to obtain the final values plotted in **Fig. S2**.

### Analysis of sequencing data

Sequencing reads were trimmed to the first 40 bases, demultiplexed and aligned against a reference database of the intended ligation products using exact matching. Genotyping data was analyzed as follows. Genotyping probe pairs were evaluated using an exact binomial test with a null model of equal abundance between the wild type probe and the mutant probe. A SNP was called if the binomial p-value was <0.001, the total reads mapped to the probe pair was >100, and the fold change between the probes was >1.5. A locus with read counts that failed any of these criteria was not called, and thus considered a “wildcard”. Each set of 20 SNP calls and wildcards is referred to as the SARS-CoV-2 genotype associated with a given sample. We utilized a network graph-based approach to visualize the relationships of genotypes detected in the Baltimore COVID-19 cohort. Genotypes were represented as nodes and were connected if there was no disagreement between the pair of genotypes (wildcards could match anything). We utilized the R package *igraph* to determine which set of samples would result in the largest number of unique genotypes (the “maximal independent vertex set”).

## Acknowledgements

This work was made possible by support from the Johns Hopkins Medicine Discovery Fund in the form of an Innovation Award, a grant from the National Cancer Institute’s Innovative Molecular Analysis Technology (IMAT) Program and an administrative supplement from the Cancer Moonshot initiative (CA202875, CA18099607), a Prostate Cancer Foundation Young Investigator Award, the JHCEIRS (contract number HHSN272201400007C) and a grant from the National Institute of Allergy and Infectious Disease to the HIV Prevention Trials Network (HPTN) Laboratory Center (AI068613). We are grateful to Beatriz Parra for providing the Zika virus isolate used in the limit of detection studies and to Priya Duggal and Rebecca Munday for assistance designing host mRNA targeting probes. Many thanks to Haiping Hao at the Johns Hopkins Transcriptomics and Deep Sequencing Core Facility for critical assistance with sequencing.

## Author Contributions

J.J.C. and J.G. performed the COVID-19 NP swab specimen experiments and assisted with data analysis. M.L.R. assisted in experimental design, probe design, and helped write the manuscript. S.H.E. helped write the manuscript. D.M. analyzed the genotyping data sets. B.S. performed the optimization experiments. A.T. and J.J.C. performed the reference organisms limit of detection studies. J.H., K.S.-S., R.E.R, A.P. and H.M. provided clinical specimens as served as technical consultants. X.Z. assisted with data analysis. K.H. designed the human splicing RASL-seq probe panel. M.S. developed software to design cRASL-seq probe sets, including the genotyping probes. H.B.L. conceived of the project and oversaw all aspects of the experimental design and data analyses.

---

**Supplemental Figure 1.**
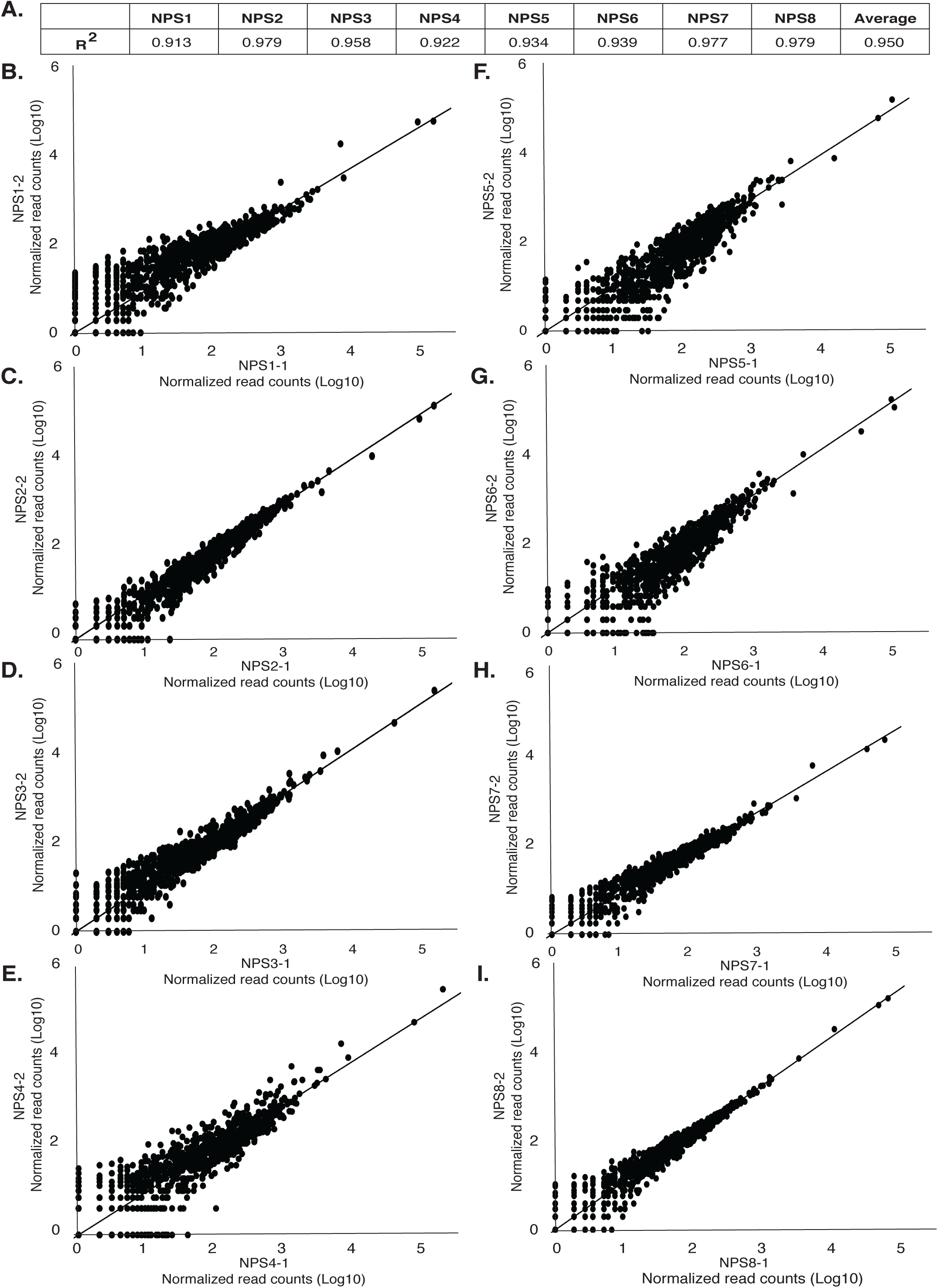
Quantification of host panel RASL-seq data. Eight NP swab specimens were subjected to RASL-seq in duplicate. Normalized read counts are plotted for each probe set.

**Supplemental Figure 2.**
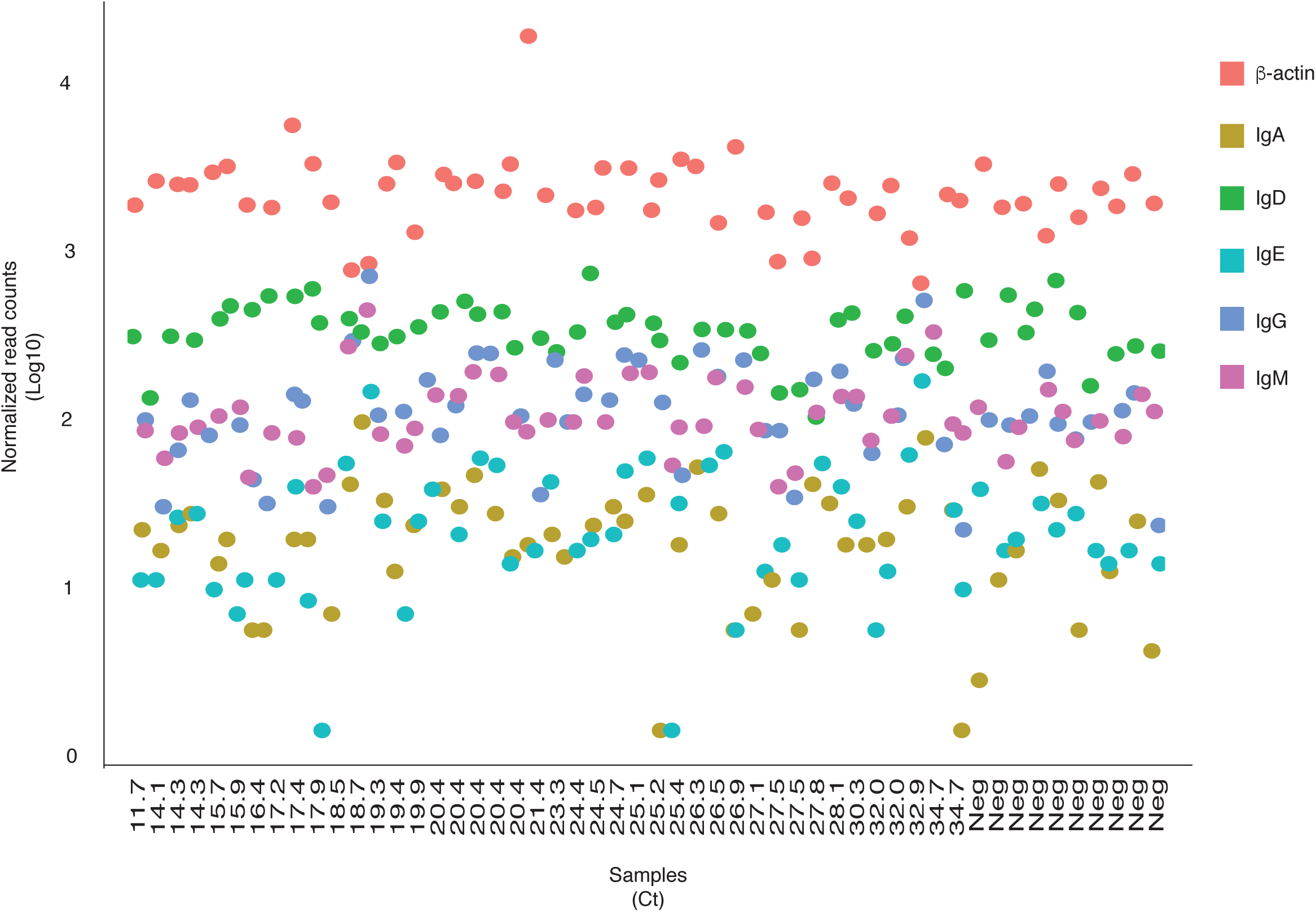
RASL-seq measurement of host immune gene expression directly from unextracted COVID-19 patient swab specimens. Results from quantification of the immunoglobulin family of genes (IgA, IgD, IgE, IgG, IgM) and a housekeeping target, β-actin, are shown. Sums of normalized read counts from all probe sets targeting the Ig or β-actin transcript are plotted (y-axis) for samples with decreasing SARS-CoV-2 viral load (increasing RT-qPCR Ct values left to right, labeled below the axis).

**Supplemental Figure 3.**
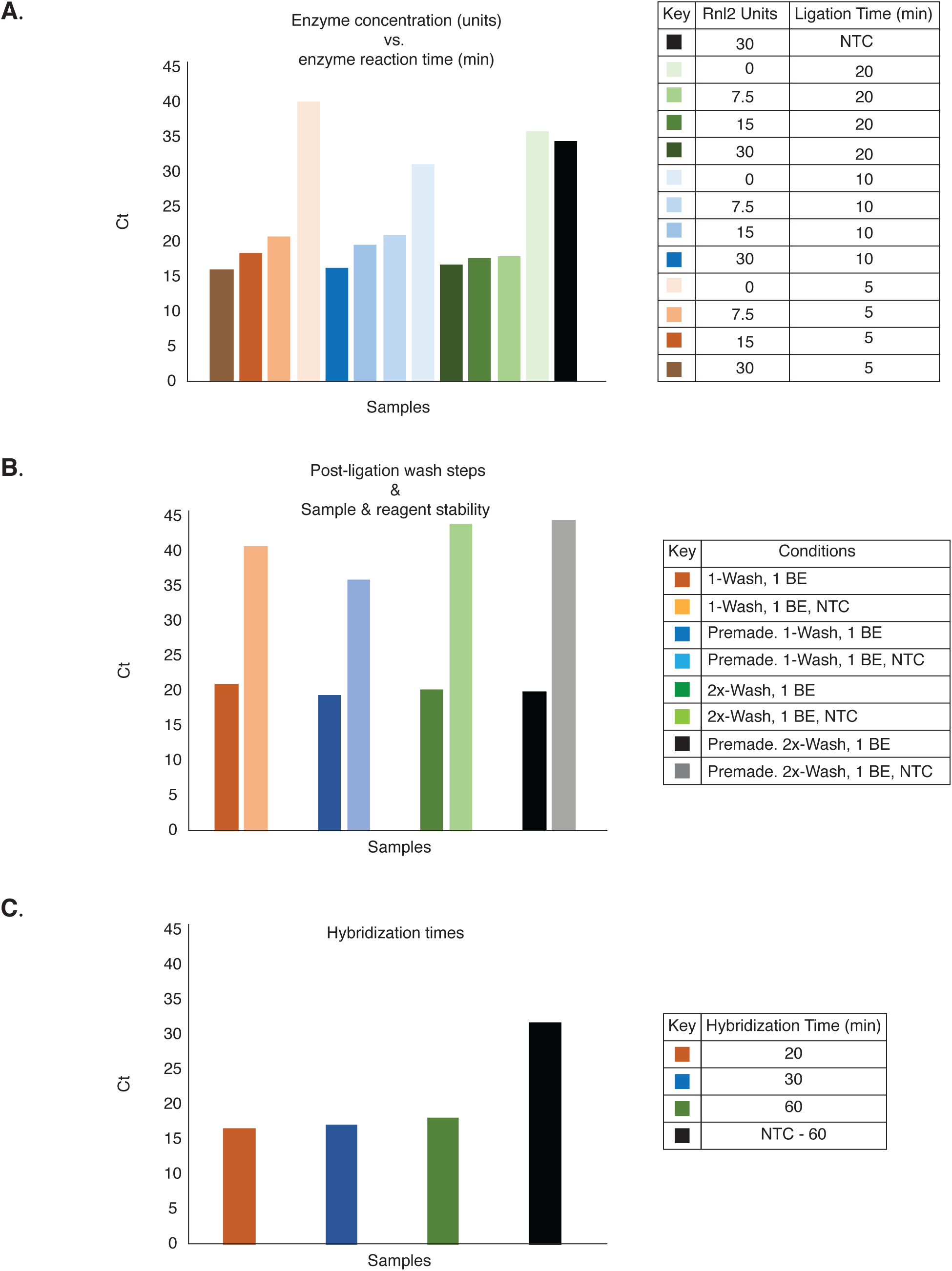
Optimization of the cRASL-seq assay. 1,000 pfu of H1N1 influenza virus collected via nasopharyngeal swab was used in standard cRASL reactions together with probe sets targeting the influenza M-segment. All reactions were analyzed by qPCR probes specific for the ligated product. **A**. Effects on RNA templated and non-templated ligation efficiency with varying Rnl2 amounts (0, 7.5, 15, and 30 units) and Rnl2 ligation times (5, 10, and 20 minutes). **B**. Effects of varying post-ligation wash steps and sample-reagent reaction storage times. **C**. Effects of varying hybridization times (20, 30, and 60 minutes). NTC, no template control. Wash, 1x-SSC. BE, buffer exchange into Rnl2 reaction buffer. Premade, complete cRASL reaction stored for 24 hours at room temperature prior to start of assay. All reactions were performed in duplicate, averages are plotted.

**Supplemental Figure 4.**
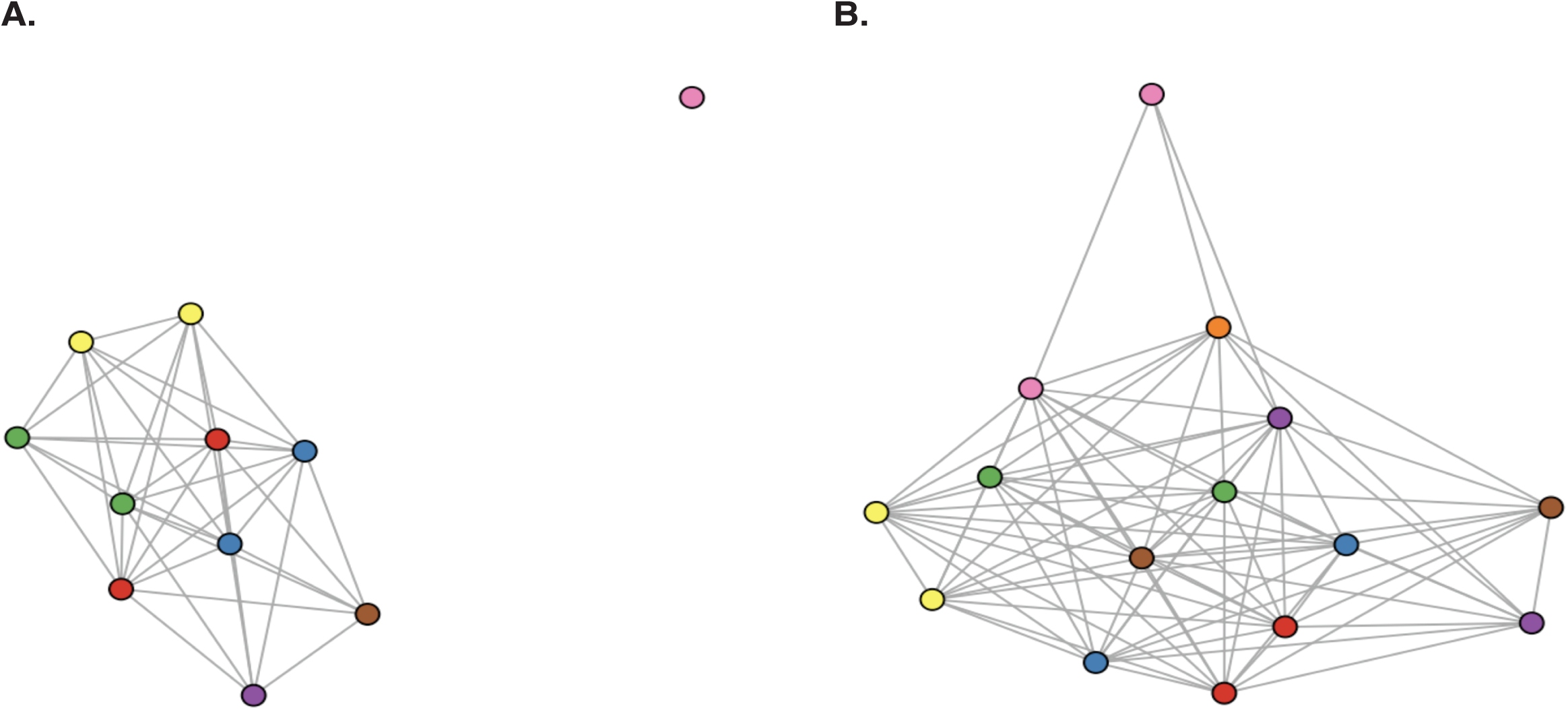
Network graph analysis of replica SNP-genotyping data. Nodes represent sample genotypes and are linked if they do not have conflicting SNPs at any position (as in Fig. 3D). Technical replicas are distinguished by color. Samples lacking sufficient called SNPs were not included (**A**. 5 SNPs to pass QC, **B**. 3 SNPs to pass QC).

**Supplemental Figure 5.**
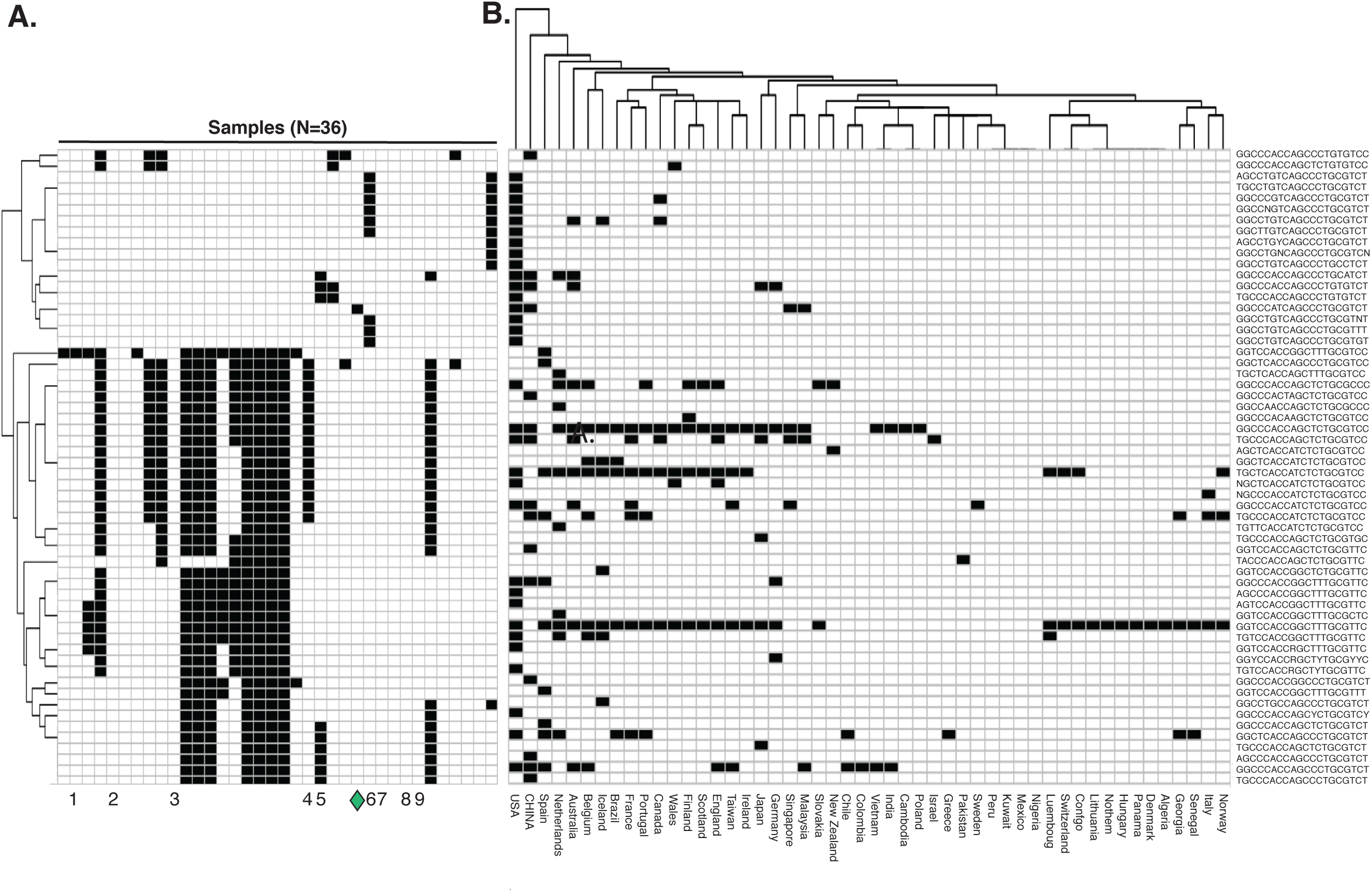
Alignment of detected SARS-CoV-2 genotypes against the GISAID database. **A**. Comparison of 20 SNP genotypes (columns) detected in Figure 3C against genotypes present in the GISAID database (rows). Rows were hierarchically clustered; column order is maintained from Fig. 3C. A match (box shaded black) means there were no conflicting SNPs between the sample and GISAID genotype. Independent genotypes (1-9) and reference isolate 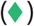 are indicated below, as in Fig. 3C. **B**. Heatmap linking GISAID genotypes (rows, same order as in **A**) with geographic location (columns), requiring exact matching to at least one isolate from the geographic region.

